# Biotransformation of D-xylose to D-xylonic acid coupled to medium chain length polyhydroxyalkanoate production in cellobiose-grown *Pseudomonas putida* EM42

**DOI:** 10.1101/702662

**Authors:** Pavel Dvořák, Jozef Kováč, Víctor de Lorenzo

## Abstract

Co-production of two or more desirable compounds from low-cost substrates by a single microbial catalyst could greatly improve the economic competitiveness of many biotechnological processes. However, reports demonstrating the adoption of such co-production strategy are still scarce. In this study, the ability of genome-edited strain *Psudomonas putida* EM42 to simultaneously valorise D-xylose and D-cellobiose -two important lignocellulosic carbohydrates -by converting them into the platform chemical D-xylonic acid and medium chain length polyhydroxyalkanoates, respectively, was investigated. Biotransformation experiments performed with *P. putida* resting cells showed that promiscuous periplasmic glucose oxidation route can efficiently generate extracellular xylonate with high yield reaching 0.97 g per g of supplied xylose. Xylose oxidation was subsequently coupled to the growth of *P. putida* with cytoplasmic β-glucosidase BglC from *Thermobifida fusca* on D-cellobiose. This disaccharide turned out to be a better co-substrate for xylose-to-xylonate biotransformation than monomeric glucose. This was because unlike glucose, cellobiose did not block oxidation of the pentose by periplasmic glucose dehydrogenase Gcd, but, similarly to glucose, it was a suitable substrate for polyhydroxyalkanoate formation in *P. putida*. Co-production of extracellular xylose-born xylonate and intracellular cellobiose-born medium chain length polyhydroxyalkanoates was established in proof-of-concept experiments with *P. putida* grown on the disaccharide. This study highlights the potential of *P. putida* EM42 as a microbial platform for the production of xylonic acid, identifies cellobiose as a new substrate for mcl-PHA production, and proposes a fresh strategy for the simultaneous valorisation of xylose and cellobiose.

## Introduction

Up to 220 million tonnes of lignocellulosic and cellulosic waste are potentially available for biotechnological purposes only in the EU every year (Searles and Malins, 2013). Lignocellulose can be decomposed by physical or chemical pre-treatment to cellulose, hemicellulose, and lignin and these fractions can be further hydrolysed enzymatically to monomeric sugars and lignin-derived aromatics serving as cheap substrates for microbial fermentations and biosynthesis of value-added chemicals (VAC) (Mosier *et al.*, 2005; Kawaguchi *et al.*, 2016). Economics of these bioprocesses is regrettably still often unsatisfactory but can be significantly improved by parallel valorisation of two or more lignocellulosic substrates. This is allowed by co-streaming of carbon from several sources into a single valued compound or by simultaneous production of two or more VAC (Dumon *et al*., 2012; Li *et al*., 2017; Larroude *et al*., 2018; Baral *et al*., 2019; Wang *et al*., 2019). Co-production of extracellular and intracellular biochemicals is desirable for facilitated downstream processing (Wang *et al.*, 2019). However, studies reporting such parallel biomanufacturing of two VAC from the second generation carbon sources are infrequent and well-defined cell factories that could efficiently perform these tasks are scarce.

The soil bacterium and growingly used robust platform strain *P. putida* KT2440 can naturally assimilate a spectrum of aromatic compounds and organic acids but only a few plant biomass-derived carbohydrates: glucose, mannose and fructose (Linger *et al.*, 2014; Belda *et al.*, 2016; Nikel and de Lorenzo, 2018; Jayakody *et al.*, 2018). Its metabolism was engineered to reach out to other sugars, including carbohydrates typically produced upon (hemi)cellulose hydrolysis or pyrolysis (Meijnen *et al.*, 2008; Linger *et al.*, 2016; Löwe *et al.*, 2018). In a recent work, *P. putida* EM42, a *P. putida* KT2440-derived strain with streamlined genome and better physiological properties (Martínez-García *et al.*, 2014), was empowered with a xylose transporter and isomerase pathway from *Escherichia coli* along with a cytoplasmic β-glucosidase BglC from *Thermobifida fusca* (Dvořák and de Lorenzo, 2018). This allowed the resulting strain to co-utilise and grow on mixtures of D-glucose, D-cellobiose, and D-xylose. However, the mix of carbohydrates was metabolized and converted into CO_2_ and biomass without any other return.

There are various possibilities to use *P. putida* for VAC biomanufacturing from glucose and cellobiose (Poblete-Castro *et al.*, 2012; Loeschcke and Thies, 2015). *P. putida* KT2440 has been traditionally employed as a model organism for the production of medium-chain-length polyhydroxyalkanoates (mcl-PHA), biodegradable polyesters applicable for manufacturing of packaging materials, textile, or medical implants (Chen, 2009; Prieto *et al.*, 2016; Li *et al*., 2017). The mcl-PHA have better elastomeric properties and broader application potential than short-chain-length PHA produced by *Cupriavidus necator* or recombinant *E. coli* (Chen, 2009). Synthesis of mcl-PHA was demonstrated from fatty acids and unrelated substrates such as acetate, ethanol, glycerol, or some sugars including glucose (Prieto *et al.*, 2016) but never from cellodextrins such as cellobiose. In a previous study, we also identified the ability of *P. putida* EM42 to oxidize D-xylose to D-xylonic acid, a platform molecule of considerable biotechnological interest (Werpy and Petersen, 2004; Toivari *et al.*, 2012; Mehtiö *et al.*, 2016; Dvořák and de Lorenzo, 2018). D-xylonate was reported to be used as a complexing agent or chelator, as a precursor of polyesters, 1,2,4-butanetriol, ethylene glycol or glycolate, and it can serve as a cheap, non-food derived alternative for D-gluconic acid (Toivari *et al.*, 2012). Xylonate is naturally formed in the first step of oxidative metabolism of xylose by some archaea, bacteria, and fungi *via* the action of D-xylose or D-glucose dehydrogenases. Production of xylonate was reported for instance in *Gluconobacter oxydans*, in several *Pseudomonas* strains including *P. fragi, P. taiwanensis or P. putida S12*, or in *Klebsiella pneumoniae (Buchert et al., 1988; Meijnen et al*., 2009; Köhler *et al*., 2015; Wang *et al.*, 2016). Several other microorganisms including *Escherichia coli* or *Saccharomyces cerevisiae* were engineered for xylonate production from xylose (Nygård *et al.*, 2011; Liu *et al.*, 2012; M. Toivari *et al.*, 2012; Gao *et al*., 2019). High production costs nonetheless hinder commercialization of both xylonate and mcl-PHA and new solutions are appealing for easing the biomanufacture of these chemicals (Chen, 2009; M. H. Toivari *et al.*, 2012; Mehtiö *et al.*, 2016, Li *et al*., 2017). Their co-production from the second generation carbon sources can thus be a promising approach in this context.

We present below our efforts to merge the advantages of *P. putida* EM42 as a natural xylonate producer with the ability of an engineered variant to grow on cellulose-derived substrate. Our results confirm that *P. putida* EM42 can convert xylose to xylonate with a high yield with its periplasmic glucose oxidative pathway and release the acid in the medium (Fig. 1). Furthermore, we show that xylonate production is inhibited in the presence of glucose but does occur in the cellobiose-grown recombinant strain. Most importantly, we demonstrate that periplasmic production and release of xylonate by cellobiose-grown *P. putida* EM42 is accompanied by parallel accumulation of mcl-PHA in the cells.

**Figure 1.**
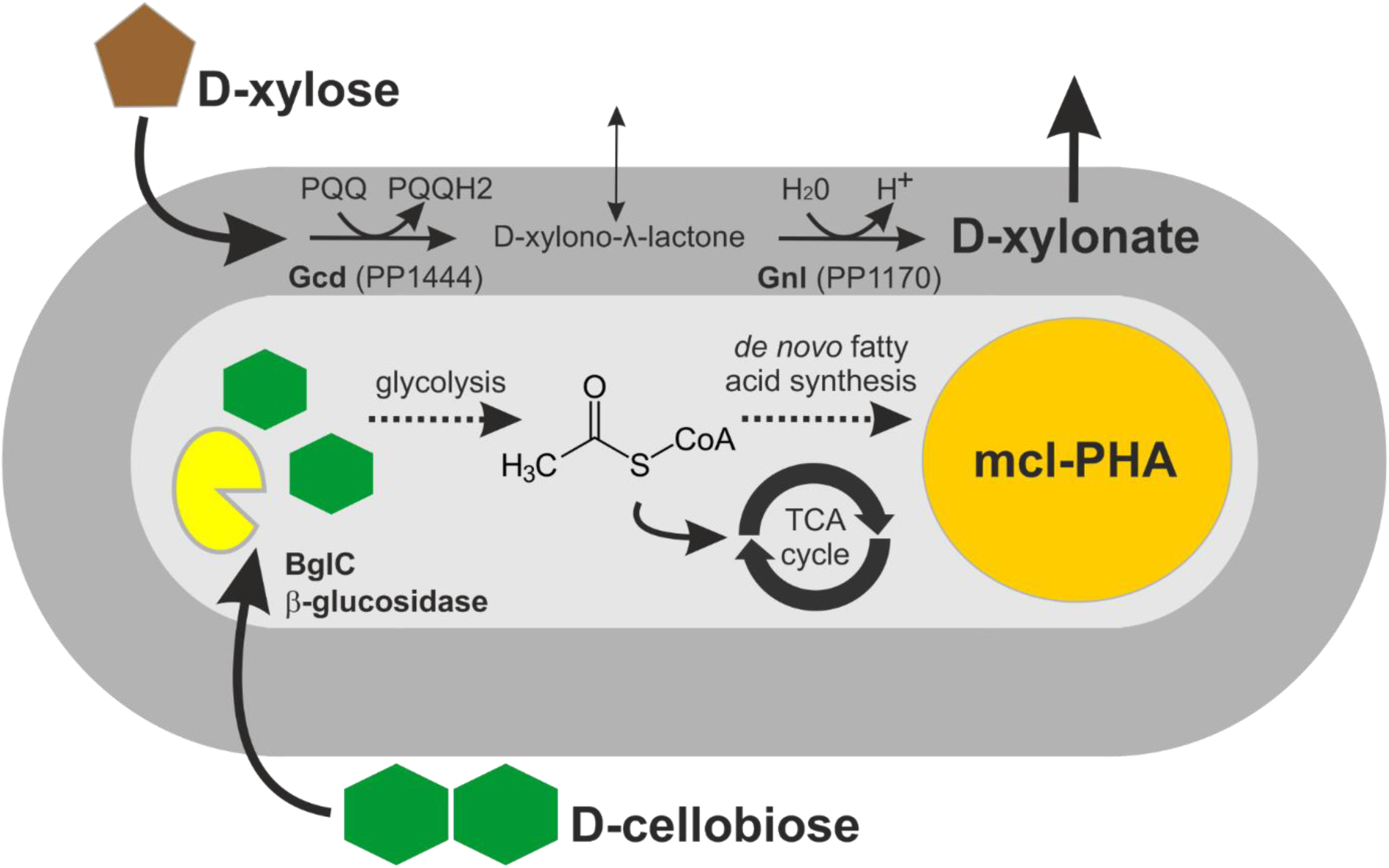
Co-production of D-xylonic acid and medium chain length polyhydroxyalkanoates from D-xylose and D-cellobiose, respectively, in *bglC*+ *P. putida* EM42. Innate periplasmic oxidative route and introduced cytoplasmic β;-glucosidase BglC from *Thermobifida fusca* allow simultaneous valorisation of D-xylose and D-cellobiose in *Pseudomonas putida* EM42. D-xylose is oxidised to platform chemical D-xylonic acid which is released into the medium. D-cellobiose, on the other hand, is transported into the cell, cleaved in two D-glucose molecules by BglC and gives rise to acetyl-CoA, a precursor molecule for the production of intracellular biopolymers (polyhydroxyalkanoates, PHA) *via de novo* fatty acid synthesis in nitrogen-limited conditions. Periplasmic space and cytoplasm are shown in dark and pale grey, respectively. Abbreviations: Gcd, glucose dehydrogenase; Gnl, gluconolactonase; PQQ and PQQH, pyrroloquinoline quinone and its reduced form, respectively; TCA cycle, tricarboxylic acid cycle; mcl-PHA, medium chain length polyhydroxyalkanoates.

## Results and Discussion

### Biotransformation of xylose to xylonate by P. putida EM42 resting cells

Periplasmic xylose conversion to xylonate was previously identified as a competing reaction for xylose assimilation by recombinant *P. putida* EM42 during a five-day cultivation experiment (Dvořák and de Lorenzo, 2018). Periplasmic glucose dehydrogenase was shown to be a crucial component for xylose oxidation in our strain as well as in several xylonate producing bacteria including *Klebsiella pneumoniae* and some other pseudomonads (Hardy *et al.*, 1993; Meijnen *et al.*, 2008; Köhler *et al*., 2014; Wang *et al.*, 2016; Dvořák and de Lorenzo, 2018). In *P. putida* KT2440, and correspondingly also in strain EM42, membrane-bound glucose dehydrogenase Gcd (PP1444) oxidises xylose to xylonolactone with pyrroloquinoline quinone (PQQ) as a cofactor. Lactone can then open spontaneously in the presence of water or might be converted to xylonate with the help of gluconolactonase Gnl (PP1170). Neither xylose nor xylonate is utilised for biomass formation (Dvořák and de Lorenzo, 2018).

Here, we initially tested whether xylose can be oxidized to xylonate in a short time interval and with a high yield by *P. putida* resting cells of defined optical density. Xylonolactone concentrations were newly determined in culture supernatants using the hydroxamate method (Lien, 1959), which allowed more precise quantification of xylonate than in our previous work (Dvořák and de Lorenzo, 2018). *P. putida* EM42 cells (strains and plasmids used in this study are listed in Supplementary Table S1), pre-cultured in lysogeny broth (LB), washed and diluted to a starting OD_600_∼ 0.5, were incubated for 48 h in M9 minimal medium with 5 g L^−1^ xylose (Fig. 2A). The yield of extracellular xylonate detected in the medium at the end of the incubation was 0.95 g per g of xylose which was 85% of the theoretical maximum 1.12 g g^−1^. Lactone accumulated in small quantities (up to 0.45 g L^−1^) in the medium during the initial phase of fast xylose conversion, but its concentration then declined to zero at the end of the experiment. Dehydrogenase activity measured with whole cells reached 2.87 ± 0.21 U per gram of dry biomass. The release of sugar acid was accompanied by a pH drop in the medium from the initial 7.00 ± 0.00 to 6.15 ± 0.04 at the end of the reaction. Neither lactone nor xylonate was detected in the identical experiment repeated with *P. putida* EM42 Δ*gcd* mutant lacking glucose dehydrogenase (Fig. 2B). These experiments confirmed the importance of Gcd for D-xylose oxidation to xylonate in *P. putida* EM42 and showed that xylonolactone intermediate is converted rapidly to xylonic acid which is released into the medium rather than utilised by the cells. In contrast, a study with *P. fragi* (the best-described pseudomonad in term of xylonate production thus far), reported slow spontaneous hydrolysis and accumulation of inhibitory xylonolactone in this bacterium during the early phases of fermentation experiments (Buchert *et al.*, 1986; Buchert and Viikari, 1988). Another well-characterized xylose-oxidizing pseudomonad, *P. taiwanensis* VLB120, uses xylonate for biomass formation (Köhler *et al*., 2014). *P. putida* thus represents an attractive addition to these strains for fast high-yield production of extracellular xylonate.

**Figure 2.**
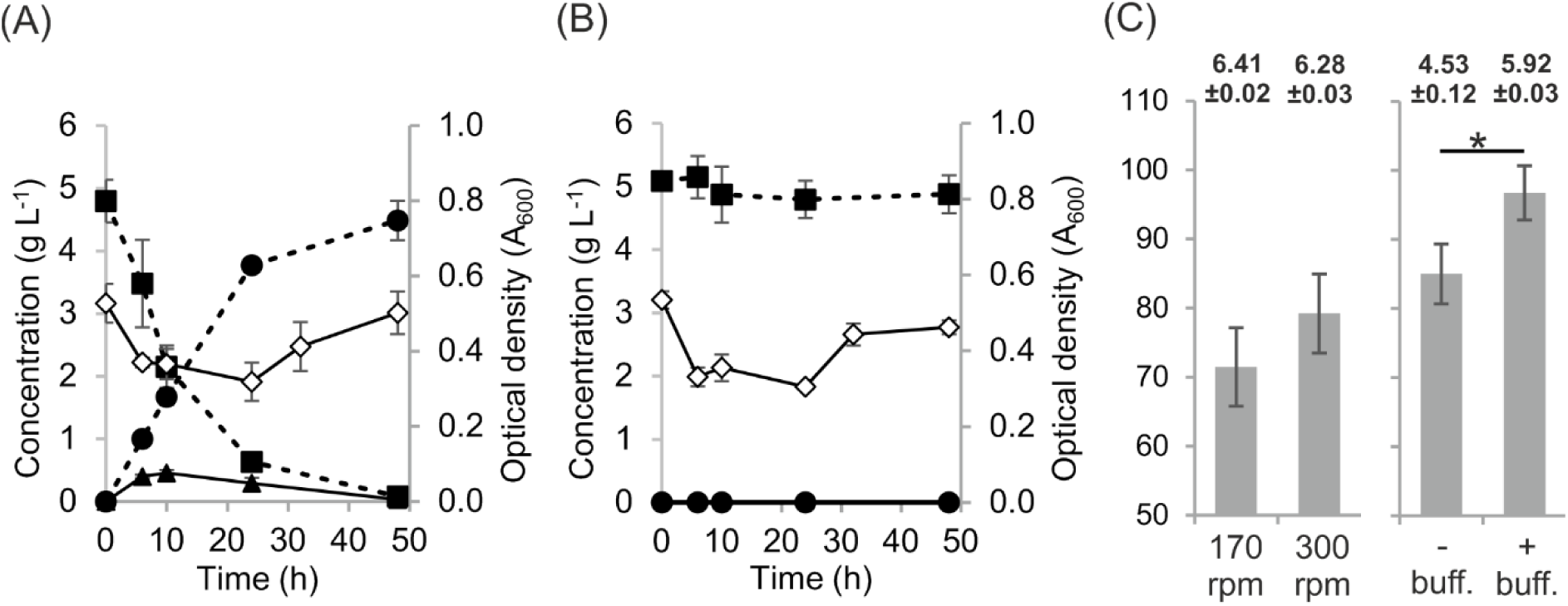
Biotransformation of xylose to xylonate by *P. putida* EM42 resting cells. Incubation of resting cells of (A) *Pseudomonas putida* EM42 and (B) its deletion mutant *P. putida* EM42 Δ*gcd* in minimal medium with 5 g L^−1^ D-xylose. Experiments were carried out in 25 mL of minimal medium in flasks shaken at 170 rpm and 30 °C. Minimal medium was inoculated to the initial A_600_ of 0.5 using cells obtained from an overnight culture in lysogeny broth. D-xylose, filled squares (▪); D-xylonate, filled circles (•); D-xylono-λ-lactone, filled triangles (▴); cell biomass, open diamonds (⋄). Data points shown as mean ± SD of three biological replicates. (C) Effect of agitation speed (left graph) and pH (right graph) on conversion of xylose to xylonic acid by *P. putida* EM42 resting cells. For evaluation of agitation speed effect, cells of A_600_ = 0.2 were incubated in medium with 5 g L^−1^ xylose with agitation of 170 or 300 rpm and the level of xylose conversion to xylonate was determined after 48 h. For evaluation of pH effect, cells of A_600_ = 1.0 were incubated in medium with or without 100 mM sodium phosphate buffer, with 10 g L^−1^ xylose and at agitation of 170, the level of xylose conversion to xylonate was determined after 48 h. The numbers above the columns show pH of the medium measured at the end of the experiment. Both experiments were carried out with resting cells collected from overnight cultures in LB medium, washed and re-suspended in shake flasks with 20 mL of M9 minimal medium. Columns represent means ± SD from at least four biological replicates from two independent experiments. Asterisk denotes significance in the difference between two means at P < 0.01 as evaluated by Student’s *t* test.

It is worth noting that the resting *P. putida* cells could be recycled and used repeatedly in five cycles of xylose oxidation to xylonic acid (Supplementary Fig. S1). The conversion reached 94 % in the first cycle, then decreased and reached 60 % in the last fifth cycle. As the optical density of the cells measured at the end of each cycle continuously decreased, the decline in productivity can be attributed mainly to the loss of the biomass in the reactions due to the centrifugation/re-suspension cycles and cell lysis (Supplementary Fig. S1). Medium pH drop detected at the end of each cycle corresponded with the level of xylose-to-xylonate conversion (Supplementary Fig. S1). This result indicates that the xylose oxidation in *P. putida* EM42 is not necessarily growth-dependent as reported with *P. fragi* (Buchert *et al.*, 1986; Buchert and Viikari, 1988). It is worth to note that a number of studies on microbial xylonate production have reported the association of xylose oxidation to a host’s growth (Toivari *et al.*, 2012; Köhler *et al.*, 2014; Wang *et al.*, 2016) but some have not. One example of the latter is recent work by Zhou and co-workers (2017) on *G. oxydans*, which could be used repeatedly for xylonate production in a bioreactor with an improved oxygen delivery system. Such cell recycling can be a promising strategy offering high xylonate yield and reduced process costs.

Since oxygen availability may become a bottleneck for the xylose-to-xylonate conversion, we next examined the effect of improved aeration through increased agitation of *P. putida* resting cells. In Zhou *et al.* (2017), the increase in agitation speed from 300 to 500 rpm enhanced the accumulation of xylonate by 25%. To check whether we could observe the same trend, we incubated resting cells in minimal medium with 5 g L^−1^ xylose at agitation of 170 or 300 rpm and the level of xylose conversion to xylonate was determined after 48 h (Fig. 2C). Xylose oxidation to xylonate was about 10% more efficient in flasks agitated at higher speed. However, the increase was only marginal.

Another variable tested was pH. Xylonate accumulation results in acidification of the medium and low pH can inhibit the activity of glucose dehydrogenase, as shown previously for *P. fragi* (Buchert *et al.*, 1986). To inspect the effect of pH, we increased the buffering capacity of the M9 medium by mixing it with 100 mM sodium phosphate buffer while escalating glucose concentration to 10 g L^−1^to intensify acidification. As shown in Fig. 2C, in these conditions EM42 cells (A_600_= 1.0) gave rise to 9.67 ± 0.39 g L^−1^ of xylonate after 48 h, while 8.50 ± 0.43 g L^−1^of xylonic acid were detected in supernatants of non-buffered cultures (Fig. 2C). The final pH determined in buffered (5.92 ± 0.03).and non-buffered (4.53 ± 0.12) cultures proved that the sodium phosphate buffer of used concentration could efficiently prevent excessive pH drop. These observations on culture conditions were considered for increasing the efficiency of xylose conversion to xylonate in subsequent experiments with growing *P. putida* cells.

### Xylose biotransformation to xylonate by P. putida EM42 growing on glucose or cellobiose

In none of the experiments mentioned above xylose oxidation to xylonate was tested during growth. Instead, the transformation experiments were preceded by the production of whole-cell catalyst biomass. Similarly to other naturally occurring or recombinant xylonate producers (Nygård *et al.*, 2011; M. Toivari *et al.*, 2012; Wang *et al.*, 2016; Zhou *et al.*, 2017; Gao *et al*., 2019) *P. putida* was grown in a medium rich in amino acids and vitamins, namely in LB (La Rosa *et al*., 2016). However, such complex media are expensive and thus unsuitable for large-scale bioprocesses. As an alternative, the growth of xylonate producing microorganism on low-cost carbon source derived *e.g.*, from lignocellulosic materials, would be desirable. D-glucose is the most abundant monomeric sugar in lignocellulosic hydrolysates prepared by using commercial enzyme cocktails with endoglucanase, exoglucanase, and β-glucosidase (Taha *et al.*, 2016) and it is also a good growth substrate for *P. putida* (del Castillo *et al.*, 2007). However, glucose is a preferred substrate for glucose dehydrogenase and might thus inhibit xylose oxidation by this enzyme. Figure 3A shows that this is exactly the case, *P. putida* EM42 cultured in minimal medium with 5 g L^−1^ glucose did not oxidise xylose during the first eight hours of the experiment, *i.e.* when glucose was consumed by the cells. As a consequence, the production of xylonate (which occurred concomitantly with growth) was postponed and less than 20 % of xylose was oxidised to the acid at the end of the two-day culture (Table 1). Inhibition of xylose transformation to xylonate by glucose was confirmed in an additional experiment using an increased concentration of the hexose (10 g L^−1^, Supplementary Fig. S2).

**Table 1.** Parameters determined in the cultures with *Pseudomonas putida* EM42 or *P. putida* EM42 pSEVA2213_*bglC* grown on D-glucose or D-cellobiose, respectively, and transforming D-xylose to D-xylonic acid. Values represent the mean ± standard deviation of three biological replicates. Parameters (except for μ) were determined after 24 h / 48 h of the culture.

| *P. putida* strain and culture conditions | $\mu$ (h<sup>-1</sup>)<sup>c</sup> | Xylonate yield (g g<sup>-1</sup> xylose) | Xylonate productivity (mg L<sup>-1</sup> h<sup>-1</sup>) | CDW (g) | pH |
| --- | --- | --- | --- | --- | --- |
| EM42 glucose | 0.58 ± 0.02 | 0.18 ± 0.02/<br>0.17 ± 0.03 | 69 ± 6/<br>32 ± 6 | 1.73 ± 0.12/<br>1.60 ± 0.06 | 6.26 ± 0.01/<br>5.81 ± 0.03 |
| *bglC*+ EM42 cellobiose | 0.27 ± 0.03 | 0.30 ± 0.06/<br>0.48 ± 0.09 | 114 ± 18/<br>93 ± 13 | 1.46 ± 0.13/<br>1.88 ± 0.05 | 6.16 ± 0.06/<br>5.19 ± 0.03 |
| *bglC*+ EM42 cellobiose optA<sup>a</sup> | 0.30 ± 0.02 | 0.35 ± 0.02/<br>0.50 ± 0.01 | 144 ± 8/<br>102 ± 3 | 2.41 ± 0.11/<br>2.18 ± 0.07 | 6.27 ± 0.08/<br>6.00 ± 0.04 |
| *bglC*+ EM42 cellobiose optB<sup>b</sup> | 0.24 ± 0.01 | 0.41 ± 0.09/<br>0.52 ± 0.08 | 156 ± 32/<br>99 ± 13 | 0.86 ± 0.02/<br>1.24 ± 0.14 | 6.24 ± 0.02/<br>5.90 ± 0.01 |

<sup>a</sup> Cultures were carried out in flasks with M9 minimal medium buffered with 100 mM sodium phosphate buffer and shaken at 300 rpm.
<sup>b</sup> Cultures were carried out in flasks with M9 minimal medium with reduced content of nitrogen, buffered with 100 mM sodium phosphate buffer and shaken at 300 rpm.
<sup>c</sup> The specific growth rate ( $\mu$ ) was determined during exponential growth. CDW, cell dry weight

**Figure 3.**
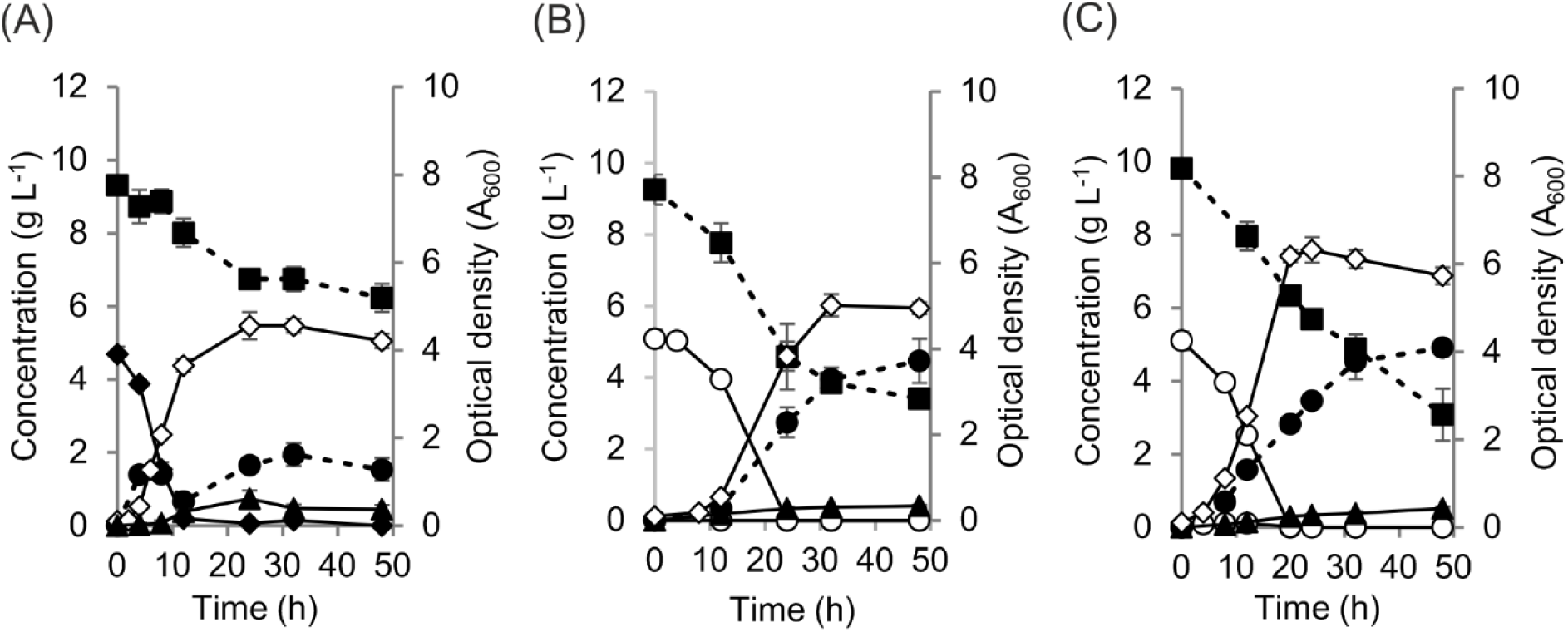
Biotransformation of D-xylose to D-xylonate by *P. putida* EM42 growing on D-glucose or D-cellobiose. Three-day cultures of (A) *Pseudomonas putida* EM42 in minimal medium with 10 g L^−1^ D-xylose and 5 g L^−1^ D-glucose used as a sole carbon source for growth. (B,C) Cultures of *Pseudomonas putida* EM42 pSEVA2213_*bglC* in minimal medium with 10 g L^−1^ D-xylose and 5 g L^−1^ D-cellobiose used as a sole carbon source. Experiments (A) and (B) carried out in 25 mL of minimal medium in flasks shaken at 170 rpm and 30 °C. Minimal medium was inoculated to the initial A_600_ of 0.1 using cells obtained from an overnight culture in lysogeny broth. Experiment (C) was performed in flask with 25 mL of minimal medium buffered with 100 mM sodium phosphate buffer and shaken at 300 rpm (30 °C). Cells used for inoculation of the main culture to the initial A_600_ of 0.1 were pre-grown overnight in minimal medium with 5 g L^−1^ D-cellobiose. D-xylose, filled squares (▪); D-xylonate, filled circles (•); D-xylono-λ-lactone, filled triangles (▴); D-glucose, filled diamonds (♦); D-cellobiose, open circles (○); cell biomass, open diamonds (⋄). Data points shown as mean ± SD of three biological replicates. Please, note that the elevated xylonate concentrations detected after 4 and 8 h in the culture (A) do not reflect the real levels of the xylose oxidation product. Hydroxamate method (Lien, 1959) used here for xylonate quantification was originally designed for the detection of gluconate, which temporarily accumulated in the culture medium during glucose utilisation in (A). Accumulation of gluconate at the times 4 and 8 h was verified also by the specific D-Gluconic Acid/D-Glucono-δ-lactone Assay Kit (Megazyme, data not shown).

We attempted to bypass this bottleneck by employing D-cellobiose as an alternative growth substrate for *P. putida*. D-cellobiose is a disaccharide composed of two β-glucose monomers linked by a β(1→4) bond. It is a by-product of cellulose saccharification with standard commercial mixtures of cellulases but becomes a predominant product when β-glucosidase is omitted from the cocktail (Chen, 2015). Well-defined microbial hosts capable of efficient cellobiose utilisation are therefore desirable because they can be applied in simultaneous saccharification and fermentation of cellulose for production of VAC while the process cost is reduced as addition of expensive β-glucosidase is not needed (Ha *et al.*, 2011; Chen, 2015; Parisutham *et al.*, 2017).

Previous work revealed that a recombinant *P. putida* EM42 derivative which expressed β-glucosidase gene *bglC* from *T. fusca* grew rapidly on D-cellobiose as a sole carbon source (Fig. 1; Dvořák and de Lorenzo, 2018). In this case, cellobiose enters *P. putida* cells through the glucose ABC transporter and it is then cleaved by BglC to two glucose molecules which are further processed in the cytoplasm. The peripheral glucose oxidative pathway probably does not play a role in cellobiose uptake. Hence, it was presumed that cellobiose could be used instead of glucose as a growth substrate for *P. putida* while xylose would be oxidised by non-occupied Gcd (Fig. 1). To test this hypothesis, we cultured *P. putida* EM42 pSEVA2213_*bglC* in minimal medium with 5 g L^−1^ cellobiose and 10 g L^−1^ xylose. Cellobiose was consumed within the initial 24 h of the culture under conditions described in the legend of Figure 3. No glucose was detected in the medium. During the same time interval, 2.75 ± 0.42 g L^−1^ of xylonate were produced from xylose with average volumetric productivity 114 mg L^−1^ h^−1^ which was 65 % higher than in the culture on glucose (Table 1). Xylose oxidation was fastest during the initial 32 h of the exponential growth phase and then slowed down in the stationary phase. Xylonate yield at the end of the two-day experiment was 0.48 ± 0.09 g g^−1^ xylose. Minor quantities of xylonolactone were detected in supernatant during the whole course of the culture (Fig. 3B). Xylonate production and cellular growth were accompanied by acidification of the medium: the pH decreased from 7.00 ± 0.00 to 6.16 ± 0.06 and 5.19 ± 0.03 after 24 and 48 h of culture, respectively.

The xylonate productivity after initial 24 h of the exponential growth further increased 1.26-fold (to 144 mg L^−1^ h^−1^) when the *bglC*^+^ *P. putida* EM42 strain was pre-grown in minimal medium with cellobiose and cultured in the modified conditions used previously with resting cells (100 mM sodium phosphate buffer and 300 rpm; Fig. 3C and Table 1). Then, the cells entered the stationary growth phase and xylonate production during the additional 24 h of culture was comparable with the former experiment with cells grown in standard M9 medium at 170 rpm. This cannot be attributed to pH because the buffering of the medium was more efficient (Table 1). We argue that suboptimal oxygen supply in shake flasks might be the limiting factor preventing efficient xylose oxidation by dense culture in the stationary period. In any case, these experiments indicate that cellobiose, an abundant cellulosic carbohydrate, does not inhibit xylose oxidation to xylonate in *P. putida* and can thus be used as a growth substrate for cells performing this biotransformation.

### Co-production of xylonate and PHA by P. putida EM42 grown on cellobiose

The ability of *P. putida* to both metabolize cellobiose in the cytoplasm and oxidize xylose by the periplasmic pathway paved the way for parallel co-production of the two biotechnologically relevant compounds – xylonate and mcl-PHA. The mcl-PHA have been reported to be co-produced with alginate oligosaccharides from glucose or glycerol (Guo *et al*., 2011; Licciardello *et al*., 2016) or with rhamnolipids from fatty acids (Hori *et al*., 2011). Also, D-xylonic acid was generated simultaneously with xylitol or bioethanol from xylose and glucose (Wiebe *et al*., 2015; Zhu *et al*., 2019). However, the synthesis of mcl-PHA along with the release of xylonate has not yet been reported. To this end, we first examined the formation of PHA granules in cellobiose-grown *P. putida* cells. As shown in Supplementary Figure S2, flow cytometry and confocal microscopy identified PHA in the bacteria (Experimental procedures and Results and discussion in Supporting information).

This simple test indicated that *P. putida* EM42 *bglC*^+^ metabolized cellobiose to the monomeric glucose, then to acetyl-CoA and next channeled this metabolic intermediate towards the formation of the polymer. In order to verify that PHA could be generated along with xylonate production, the *bglC*^+^ strain was pre-cultured in nitrogen-rich LB medium (to avoid any PHA accumulation) and then grown in nitrogen-limited M9 medium with 100 mM sodium phosphate buffer, 5 g L^−1^ cellobiose, and 10 g L^−1^ xylose (Fig. 4). Sugar and xylonic acid concentrations were determined in culture supernatants while intracellular PHA formation was followed by flow cytometry and confocal microscopy. As shown in Figures 4A, 4B, and 4C, xylonate and PHA were produced simultaneously during the initial 48 h of the three-day experiment. Cellular polymer content increased during the first two days and then declined towards the end of the experiment (Figs. 4B and 4C). This trend correlated with the presence of the carbon source (cellobiose and glucose) in the medium (Fig. 4A). As in previous experiments, cellobiose was almost completely consumed within the initial 24 h. However, uptake of the disaccharide was this time accompanied by the appearance of glucose in the medium, which reached its maximum concentration (1.61 ± 0.37 g L^−1^) at 12 h of the culture. Under these circumstances, it became apparent that the secreted glucose affected xylose oxidation by Gcd; only ∼ 25 % of the pentose was converted to xylonate at the end of the experiment. Although we do not have a trivial explanation for such unexpected release of glucose, we speculate that it could be due to [i] slower growth (μ = 0.19±0.01 h^−1^) under nitrogen limitation as compared to the standard M9 medium (μ = 0.30±0.02 h^−1^; Fig. 3C) and/or [ii] an imbalance between the knocked-in BglC β-glucosidase and the innate Glk glucokinase (PP1011) activities stemming from difference in composition of pre-culture (LB) and culture (M9 with cellobiose) medium (see a scrutiny of these possibilities in Results and discussion, Experimental procedures, and Fig. S4 in Supporting information).

**Figure 4.**
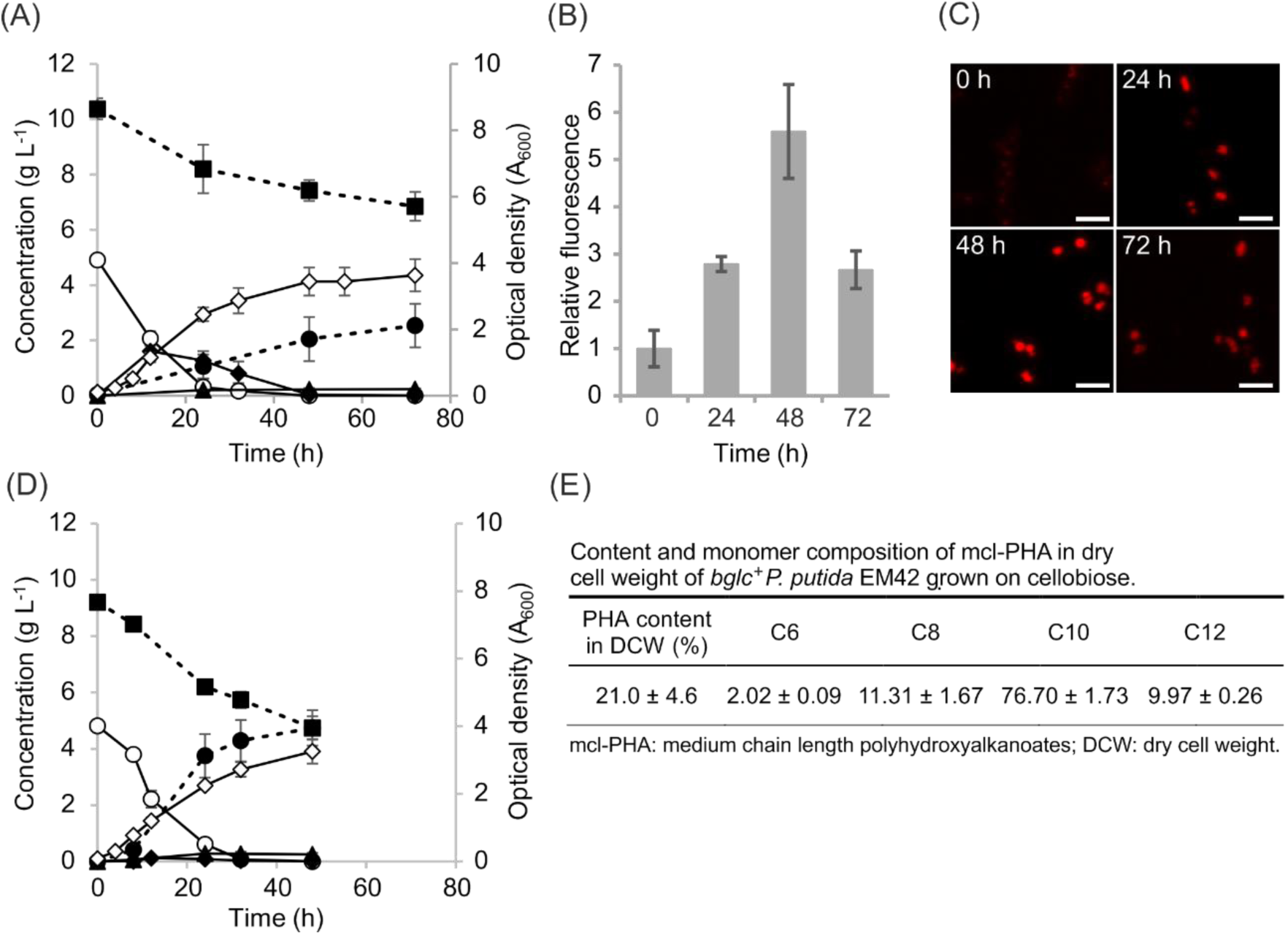
Co-production of D-xylonic acid and PHA from D-xylose and D-cellobiose, respectively, by cellobiose grown *P. putida* EM42 pSEVA2213_*bglC*. (A) Initial culture inoculated from overnight pre-culture in lysogeny broth was carried out in 25 mL of nitrogen-limited M9 minimal medium with 100 mM sodium phosphate buffer, 5 g L^−1^ cellobiose, and 10 g L^−1^ xylose in flasks shaken at 300 rpm and 30 °C. (B) Relative fluorescence of bacterial population analysed by flow cytometry every 24 h during the three-day culture. Cells were stained by Nile Red and processed as described in Supplementary Information. (C) Confocal microscopy of *P. putida* cells collected at denoted time intervals. Stained bacteria were processed as described in Supplementary Information. White scale bars show 2 μm distance. (D) Culture inoculated from overnight pre-cultures in M9 minimal medium with 5 g L^−1^ cellobiose was carried out in the same conditions as were described for (A). (E) Content and monomer composition of medium chain length polyhydroxyalkanoates in dry cell weight of *P. putida* EM42 pSEVA2213_*bglC* cells collected at the end of the two-day culture. D-xylose, filled squares (■); D-xylonate, filled circles (●); D-xylono-λ-lactone, filled triangles (▴); D-glucose, filled diamonds (♦); D-cellobiose, open circles (○); cell biomass, open diamonds (⋄). Data points and columns in (A), (B), and (C) show mean ± SD of three biological replicates.

To overcome this bottleneck, *P. putida* EM42 *bglC*^*+*^ cells were pre-grown overnight in standard M9 medium with 5 g L^−1^ cellobiose instead of LB. Faster growth of the main cultures (μ = 0.24±0.01 h^−1^) in the nitrogen-limited M9 medium with cellobiose and xylose was then indeed observed and only minute concentrations of glucose (up to 0.12 g L^−1^) were detected in the supernatants during first 24 h (Fig. 4D). As a consequence, the volumetric productivity of xylonate during this period increased 3.5-fold (from 44±18 mg L^−1^ h^−1^ to 156±32 mg L^−1^ h^−1^) when compared with the previous experiment shown in Fig. 4A. Xylonate yield was 2.4-times higher and reached 0.52±0.08 g g^−1^ xylose after 48 h of the culture (Table 1). Interestingly, the xylonate yield per gram of cell dry weight was 1.7-fold higher compared to the cells growing faster and reaching higher OD_600_ in M9 medium with standard nitrogen content (Fig. 3C, Table 1).

The same cultures were stopped after 48 h to quantify also PHA content within the cells which turned out to be 21 % (w/w) of cell dry weight. The biopolymer yield was 0.05 ± 0.01 g g^−1^ cellobiose. These values are close to those reported for *P. putida* KT2440 grown on glucose (Huijberts *et al*., 1992; Poblete-Castro *et al*., 2013). The yield of PHA per 1 L of the cell culture in the shake flask reached 0.26 ± 0.03 g. The monomer composition of the analysed biopolymer was also consistent with the previous reports on mcl-PHA production from glucose (Fig. 4E). The major fraction (> 75 %) was formed by 3-hydroxydecanoate, followed by 3-hydroxyoctanoate, 3-hydroxydodecanoate, and small amount of 3-hydroxyhexanoate. Taken together, the above experiments confirmed the co-production of two value-added molecules (xylonate and mcl-PHA) out of xylose and cellobiose in *P. putida*.

## Conclusion

In this work, we have exploited the metabolic versatility of *P. putida* EM42, a robust derivative of *P. putida* KT2440, for prototyping the simultaneous conversion of xylose and cellobiose into xylonate and mcl-PHA. Periplasmic oxidation of D-xylose to D-xylonic acid was first assayed with recyclable *P. putida* EM42 resting cells. Rapid transformation of pentose into free xylonate with only minor accumulation of xylonolactone intermediate was observed. Such extracytoplasmic production and secretion are advantageous over intracellular xylose oxidation: cytoplasm acidification is avoided, the reaction of interest does not cross-interfere with the host’s metabolism, and xylonate can be purified directly from the culture medium (Wang *et al.*, 2016).

We then demonstrated that xylose conversion to xylonate can be efficiently catalysed also by recombinant *P. putida* EM42 *bglC*^*+*^ growing on D-cellobiose. In contrast to monomeric glucose, which is a preferred substrate for glucose dehydrogenase in *P. putida*, the disaccharide did not compete with xylose for Gcd and was a better carbon source for growth-associated xylonate production. Importantly, cellobiose-grown *P. putida* was able to stream the carbon from disaccharide into the intracellular mcl-PHA and concomitantly oxidize xylose to xylonate. Both xylonate and PHA yields could be further increased not only through bioprocess design but also by additional genetic interventions in the host that are known to improve the two bioproductions separately. This includes *e.g*., overexpression of genes encoding the periplasmic oxidative pathway (Yu *et al*. 2018) and/or overexpression of pyruvate dehydrogenase gene *acoA* (Borrero-de Acuña *et al*., 2014). These optimisation efforts will be the subject of our further work. In any case, bioprocesses based on microbial hosts capable of parallel production of two or more VAC from cheap abundant substrates are drawing considerable attention (Dumon *et al*., 2012; Li *et al*., 2017; Larroude *et al*., 2018; Baral *et al*., 2019; Wang *et al*., 2019). We argue that the strategy shown here on example of recombinant *P. putida* EM42 expressing cytoplasmic β-glucosidase represents a promising route for valorisation of (hemi)cellulosic residues and an alternative to the xylonate and mcl-PHA bioproductions reported thus far.

## Acknowledgments

We thank Prof. Ivana Márová and Assoc. Prof. Stanislav Obruča from Brno University of Technology for the valuable discussions and provided know-how and infrastructure for PHA quantification and characterization. This work was funded by the SETH Project of the Spanish Ministry of Science (RTI 2018-095584-B-C42), MADONNA (H2020-FET-OPEN-RIA-2017-1-766975), BioRoboost (H2020-NMBP-BIO-CSA-2018), and SYNBIO4FLAV (H2020-NMBP/0500), contracts of the European Union and the S2017/BMD-3691 InGEMICS-CM funded by the Comunidad de Madrid (European Structural and Investment Funds) as well as by the Czech Science Foundation (19-06511Y).

