## Supporting information for "Biotransformation of D-xylose to D-xylonic acid coupled to medium chain length polyhydroxyalkanoate production in cellobiose-grown *Pseudomonas putida* EM42"

by

Pavel Dvořák[1]*, Jozef Kováč[1], and Víctor de Lorenzo[2]

[1]*Department of Experimental Biology (Section of Microbiology), Faculty of Science, Masaryk University, Kamenice 753/5, 62500, Brno, Czech Republic.*

[2]*Systems and Synthetic Biology Program, Centro Nacional de Biotecnología CNB-CSIC, Cantoblanco, Darwin 3, 28049 Madrid, Spain.*

* Corresponding author: Dr. Pavel Dvořák
Department of Experimental Biology (Section of Microbiology), Faculty of Science
Masaryk University, Kamenice 735/5
Brno 62500, Czech Republic


### Contents

**Supplementary experimental procedures** ……………………………………….…………3

**Supplementary results and discussion** …………………………………………………..8

**Supplementary tables** ……………………………………………………………………11

**Table S1.** Strains and plasmids used in this study.

**Supplementary figures** ……………………………………………...………………12

**Figure S1.** Recycling of *P. putida* EM42 resting cells converting D-xylose to D-xylonic acid.

**Figure S2.** Two-day culture of *P. putida* EM42 in minimal medium with 10 g $L^{-1}$ D-xylose and 10 g $L^{-1}$ D-glucose used as a sole carbon source for growth.

**Figure S3.** Polyhydroxyalkanoate (PHA) accumulation in *P. putida* EM42 cells grown on diverse carbon sources including D-cellobiose.

**Figure S4.** Activities of β–glucosidase BglC and glucokinase Glk measured in cell-free extracts prepared from $bglC^{+}$ strain of *P. putida* EM42 grown in several different conditions.

**References** …………………………………………………………...…………………16

### Supplementary experimental procedures

#### *Media and culture conditions*

Bacterial strains and plasmids used in this study are listed in Table S1. Unless stated otherwise in figure legend, overnight pre-cultures were prepared by inoculating cells directly from glycerol stock into 10 mL of lysogeny broth (LB; 10 g $L^{-1}$ tryptone, 5 g $L^{-1}$ yeast extract, 5 g $L^{-1}$ NaCl) and cells were cultured with agitation (300 rpm; Heidolph Unimax 1010 and Heidolph Incubator 1000; Heidolph Instruments, Germany) at 30 °C (*Pseudomonas putida* EM42) or at 37 °C (*Escherichia coli* Dh5α). In the morning, cells in pre-cultures were centrifuged (4,000 g, RT, 5 min), washed with M9 minimal medium (per 1 L: 7.0 g $Na_2HPO_4$ $7H_2O$, 3 g $KH_2PO_4$, 0.5 g NaCl, 1 g $NH_4Cl$) containing 2 mM $MgSO_4$ and added with 3 mL $L^{-1}$ trace element solution (Abril *et al.*, 1989). The amount of $NH_4Cl$ was reduced to 0.2 g per 1 L in experiments for the determination of PHA content in cells. Thiamine HCl (1 mM) was added to the minimal medium for cultures with *E. coli* Dh5α. Cells were re-suspended to starting OD, defined in figure legends, in Erlenmeyer flasks with the same medium containing sugar(s). All media with cells bearing pSEVA2213 plasmid contained kanamycin (50 μg $mL^{-1}$).

#### *Analytical techniques*

Changes in $A_{600}$ were followed by VIS spectrophotometry using UV/Vis spectrophotometer Ultrospec 2100 (Biochrom, UK). Concentrations of D-xylose and D-glucose in defined time intervals were determined by Xylose Assay Kit (Megazyme, Ireland) and Glucose (GO) Assay Kit (Sigma Aldrich, USA), respectively, according to the manufacturer's instructions. D-cellobiose was quantified by HPLC-LS system 920LC with a light scattering (PL-ELS) detector (Agilent Technologies, USA) equipped with Microsorb MV NH2 column (5 μm, 250 mm x 4.6 mm). MilliQ $H_2O$ and acetonitrile were used as eluents at a flow rate of 1 mL $min^{-1}$. The column temperature was 30 °C. Concentrations of D-xylonate and D-xylono-λ-lactone were determined by the hydroxamate method introduced by Lien (Lien, 1959). Samples were prepared as described previously (Dvořák and de Lorenzo, 2018) with some modifications. Briefly, 75 μL of culture supernatants were mixed with 1.3 M HCl (1:1) and heated at 100 °C for 20 min to accelerate conversion of all present xylono-γ-lactone to xylonate (heating step was omitted in xylono-γ-lactone determinations). Samples were cooled down on ice and 50 μL was added to 100 μL of hydroxylamine reagent freshly prepared by mixing 2 M hydroxylamine HCl with 2 M NaOH. After 2 min interval, 65 μL of 3.2 M HCl and subsequently 50 μL of

$FeCl_3$ solution (10 g in 100 mL of 0.1 M HCl) were added and absorbance (550 nm) measured using Victor$^2$ 1420 Multilabel Counter (Perkin Elmer, USA). Xylonate and xylono-γ-lactone concentrations were quantified with a standard curve prepared using the pure compounds (Sigma-Aldrich, USA). As the protocol with included heating step determines the mixture of both xylonate and xylono-γ-lactone, final xylonate concentration at the given time interval was calculated by subtracting xylono-γ-lactone concentration from the obtained value.

*Determination of dry cell weight and growth parameters*

Biomass was determined as dry cell weight. Samples of cultures grown in M9 minimal medium with 5 g $L^{-1}$ glucose were transferred into 2 mL pre-dried and pre-weighed Eppendorf tubes and centrifuged at 13,000 g for 10 min. The pellets were washed twice with distilled water and dried at 80 °C for 48 h. One $A_{600}$ unit is equivalent to 0.38 g $L^{-1}$ of dry cell weight based on the prepared standard curve. Specific growth rate (μ) was calculated for the exponential growth phase as a slope of the data points obtained by plotting the natural logarithm of $A_{600}$ values against time.

*Qualitative determination of PHA formation in P. putida EM42 recombinant grown on cellobiose by flow cytometry and confocal microscopy*

Bacteria were harvested from cultures and stained with Nile Red as described elsewhere (Martínez-García, Aparicio, *et al.*, 2014a). At least 25,000 cells were analysed in each sample using MACSQuant VYB cytometer (Miltenyi Biotec, Germany). For excitation, an Ar laser (543 nm, diode-pumped solid state) was used and the fluorescence of Nile red was detected at 598 nm using a 614/50 nm band-pass filter. FlowJo v.10 software (FlowJo, USA) was used for data processing. For the purpose of microscopy, cells were washed with ice-cold phosphate buffer saline (PBS; per 1 L: 8 g NaCl, 0.2 g KCl, 1.44 g $Na_2HPO_4$, 0.24 g $KH_2PO_4$, pH adjusted to 7.4 with HCl) after staining with Nile Red and finally re-suspended in 1 mL of the same buffer. Cell suspension (5 μL) was dropped on poly-L-lysine coated glass slides (Sigma-Aldrich, USA), bacteria were mounted for 60 min, covered with 5 μL of ProLong Antifade (Thermo Fisher Scientific, USA) and the slides were analysed using confocal multispectral microscope Leica TCS SP5 (Leica Microsystems, Germany).

*PHA quantification and characterization*

*P. putida* EM42 pSEVA2213_*bglC* cells were grown overnight at 30 °C with shaking (300 rpm) in 10 mL of M9 medium with kanamycin (50 µg mL$^{-1}$) and 5 g L$^{-1}$ cellobiose as a sole carbon source. Night culture was used to inoculate 50 mL of M9 medium with reduced concentration of nitrogen, kanamycin (50 µg mL$^{-1}$), 100 mM sodium phosphate buffer, and 10 g L$^{-1}$ and 5 g L$^{-1}$ of xylose and cellobiose, respectively, to the starting $A_{600}$ of 0.1. Cells were cultured at 30 °C with shaking (300 rpm). All cells were centrifuged (10,595 *g*, 4 min, 4 °C) after 48 h of cultivation and the wet weight of biomass was measured. PHA content and monomer composition in cell biomass dried for 48 h at 80 °C was determined using gas chromatography (GC) of the methanolysed polyester. Dried cell biomass (10 mg) was mixed with 0.8 ml of esterification solution containing 15 % $H_2SO_4$ in methanol with benzoic acid (5 g L$^{-1}$) as internal standard and with 1 mL of chloroform in glass vial with crimp top. Vials were incubated at 94 °C for 3h, then cooled down and the content was mixed with 0.5 mL of 50 mM NaOH. Organic phase (50 µl) with resulting methyl esters was diluted 20-times with chloroform to a final volume of 1 mL and the samples were analysed by GC system Trace 1300 with FID detector equipped with TG-WAX MS column (30 m x 0.32 mm x 0.5 µm; Thermo Fisher Scientific). Briefly, 1 µL of the diluted sample was injected into the gas chromatograph at a split ratio of 1:25 and split flow 50 mL min$^{-1}$. The temperature of the injector and FID detector was 230 °C and 240 °C, respectively. The flow rate of the nitrogen used as a carrier gas was 2 mL min$^{-1}$. The temperature ramp was as follows: 80 °C hold for 1.5 min, then increase to 150 °C at a rate of 20 °C min$^{-1}$, hold for 2.5 min, then increase to 200 °C at a rate of 30 °C min$^{-1}$, hold for 5 min, and final increase to 230 °C at a rate of 30 °C min$^{-1}$, and the temperature was held for 2 min. The resulting methyl esters were identified and quantified using calibration curves prepared with pure standards (3-hydroxyhexanoate, 3-hydroxyoctanoate, 3-hydroxydecanoate, 3-hydroxydodecanoate), which were manipulated and analysed correspondingly, and the relative molar fraction (%) of C6, C8, C10, and C12 monomers in mcl-PHA produced by *P. putida* grown on cellobiose was calculated. The mcl-PHA content (%) in cell dry weight was determined. The experiment was performed in three biological replicates.

*Determination of β-glucosidase and glucokinase activities in P. putida recombinant grown in several different conditions*

Cell-free extracts (CFE) for enzyme assays were prepared from *P. putida* EM42 pSEVA2213_*bglC* pre-grown overnight in shake flasks (300 rpm) in 10 mL of M9 medium with kanamycin and cellobiose (5 g $L^{-1}$) or in 10 mL of LB medium with kanamycin. Pre-culture in M9 medium was used for inoculation of 25 mL of M9 medium with reduced concentration of nitrogen, kanamycin, and cellobiose (5 g $L^{-1}$) to the starting $A_{600}$ of 0.1. Culture grown overnight in LB medium was inoculated in 25 mL of LB medium with kanamycin or in 25 mL of M9 medium with reduced concentration of nitrogen, kanamycin, and 5 g $L^{-1}$ of cellobiose, in both cases to the starting $A_{600}$ of 0.1. Cells were cultured at 30 °C with shaking (300 rpm). When $A_{600}$ reached 0.5, cultures were stopped by placing the flasks on ice. Cells were then centrifuged at 10,595 g for 15 minutes at 4 °C and pellets were washed with 12 mL of ice-cold PBS buffer (80 g $L^{-1}$ NaCl, 2 g $L^{-1}$ KCl, 14.4 g $L^{-1}$ $Na_2HPO_4$, 2.4 g $L^{-1}$ $KH_2PO_4$) and then frozen at -80 °C for further use. CFEs were prepared from melted cell pellets using B-PER Bacterial Protein Extraction Kit with lysozyme and DNase I (Thermo Fisher Scientific) according to the manufacturer’s protocol. The concentration of total protein in CFEs was determined using Bradford Reagent (Sigma-Aldrich/Merck).

*β-glucosidase activity*

Enzymatic assays were made with *p*-nitrophenyl β-D-glucopyranoside substrate (Sigma Aldrich/Merck). Activity was measured in 0.58 mL of 100 mM sodium phosphate buffer (pH 7.0) with 5 mM *p*-nitrophenyl β-D-glucopyranoside. Mixture was first incubated at 30 °C for 10 minutes and the reaction was then started by the addition of 20 μL of diluted CFE (with concentration of total protein of 0,01 mg $mL^{-1}$) to the reaction mixture. Samples (0.12 mL) were taken from the reaction mixture every 5 min during the time interval of 15 min. Reaction was stopped by mixing the sample with 80 μL of 1 M $Na_2CO_3$ in a well of a microtiter plate. Increasing concentration of the reaction product *p*-nitrophenol was measured spectrophotometrically at 405 nm by plate reader Infinite M200 Pro (Tecan). Specific activity was calculated as an increase of product concentration in μmol $min^{-1}$ $mg^{-1}$ of the total protein in CFE using the calibration curve prepared with pure *p*-nitrophenol (Sigma Aldrich/Merck). One unit of enzyme activity (U) corresponds to 1 μmol of *p*-nitrophenol produced per minute.

*Glucokinase activity*

Glk was assayed in microtiter plate format. Total volume of each reaction mix (200 μL) contained 40 mM Tris-HCl buffer (pH 8.2), 1 U $mL^{-1}$ of glucose-6-P-dehydrogenase, 10 mM ATP, 4 mM $MgCl_2$, 50 mM D-glucose, 4 μL of CFE and water. The reaction was started by the addition of 0.6 mM (final concentration in the reaction mix) $NADP^+$. Increasing concentration of reaction product (NADPH) was measured spectrophotometrically at 340 nm and 30 °C using plate reader Infinite M200 Pro (Tecan). Specific activity was calculated as an increase of product concentration in μmol $mL^{-1}$ $mg^{-1}$ of the total protein in CFE using the molar extinction coefficient 6.22 mM $cm^{-1}$ for NADPH. One unit of enzyme activity (U) corresponds to 1 μmol of NADPH produced per minute.

### Supplementary results and discussion

*Qualitative determination of PHA formation in P. putida EM42 recombinant grown on cellobiose*

PHA accumulate in *P. putida* and some other bacteria in form of granules which can be stained with Nile Red and detected in cells by flow-cytometry (Spiekermann *et al.*, 1999; Tyo *et al.*, 2006). This method allows for fast verification of substrate streaming toward the industrially relevant compound (Linger *et al.*, 2014). We followed this procedure to qualitatively determine the formation of PHA in EM42 pSEVA2213_*bglC* recombinant grown for 48 h in nitrogen-limited minimal medium with an excess of cellobiose (15 g $L^{-1}$). *P. putida* EM42 pSEVA2213 and *E. coli* Dh5α pSEVA2213 grown on 20 mM octanoic acid (~2.9 g $L^{-1}$) and 30 g $L^{-1}$ glucose, respectively, were used as a positive and negative controls for PHA formation. Under given conditions, PHA was detected in 63.0 ± 9.0 % of cells cultured on cellobiose (Fig. S3A). PHA positive fraction of *P. putida* pSEVA2213 cells grown on 30 g $L^{-1}$ glucose was 75.6 ± 4.6 %. Smaller fraction of PHA-positive cells and about 30 % lower median fluorescence of the bacteria metabolizing cellobiose compared to glucose (Fig. S3B) might be attributed to the slower uptake of disaccharide (Dvořák and de Lorenzo, 2018).

*Deciphering glucose accumulation during growth of bglC$^+$ EM42 cells on cellobiose in M9 minimal medium with reduced nitrogen content*

We argued that the accumulation and secretion of glucose out of the cells grown on cellobiose in M9 minimal medium with reduced nitrogen content could be caused by: (i) slower growth of bacteria in these conditions during early exponential phase and (ii) possible imbalance between heterologous BglC β-glucosidase and innate Glk glucokinase (PP1011) activities stemming from difference in composition of pre-culture (LB) and culture (M9 with cellobiose) medium. The cells grew about 35 % slower in nitrogen-limited medium ($\mu$ = 0.19±0.01 $h^{-1}$, Fig. 4A) when compared with the former cultivations in standard M9 medium ($\mu$ = 0.30±0.02 $h^{-1}$, Fig. S2) but the amount of cellobiose they consumed during the initial 12 h of the culture was almost identical (2.84±0.28 g $L^{-1}$ and 2.58±0.09 g $L^{-1}$, respectively). Hence, rapid uptake and intracellular hydrolysis of cellobiose by BglC in cultures with nitrogen-limited medium was probably not compensated by equally fast phosphorylation of resulting glucose molecules by Glk. Accumulated non-phosphorylated (thus less polar) glucose can leave the cells *e.g.*, by reverse facilitated diffusion, and compete with xylose for Gcd in the periplasm.

The imbalance between BglC and Glk activities could be exacerbated by the fact that the *bglC* gene was expressed continuously in EM42 recombinant from constitutive EM7 promoter also during pre-culture in LB medium. On the other hand, expression of *glk*, forming part of *edd*/*glk*/*gltR-II* operon, depends on native *P. putida* regulatory mechanisms and is enhanced in presence of glucose metabolism intermediates that occurred in cells only later during the main culture on cellobiose (Udaondo *et al.*, 2018). It is known that even LB contains low a concentration of free glucose (app. 0.3 mM; La Rosa *et al*., 2016) and glucose bound in polysaccharides (up to 15 mM; Molina *et al*., 2019). However, recent work of Molina and co-workers (2019) indicates that glucose in LB is not metabolized by *P. putida* but rather used as a source of electrons for electron transport chain during oxidation of hexose to gluconate and 2-ketogluconate in periplasm. These oxidized forms are mostly released in the medium, and glycolysis (including Glk) is probably not fully induced during growth in LB.

We decided to verify the effect of pre-culture and culture medium on BglC and Glk activities in *bglC*$^{+}$ *P. putida* EM42 strain. Figure S4 shows that the activity of both enzymes was substantially lower in cells pre-cultured and cultured in LB (LB-LB) than in cells pre-cultured in LB and then cultured in M9 medium with reduced nitrogen content (LB-M9L). The difference between these two conditions had a significantly higher effect on Glk activity (4.8-times higher in LB-M9L than in LB-LB) than on BglC activity (2.4-times higher in LB-M9L than in LB-LB). This difference even increased when the cells were pre-cultured in M9 medium with cellobiose and then cultured in nitrogen-limited M9 with the same carbon source. In this condition, BglC activity in cells was still 2.4-times higher when compared with LB-LB culture, while Glk activity increased 6.0-times.

Obtained activity values are comparable with former studies (Dvořák and de Lorenzo, 2018; Sánchez-Pascuala *et al*., 2019) and reveal that glucokinase is a bottleneck enzyme in the BglC-Glk pair. The results show, at the same time, that Glk activity depends more than the activity of BglC on pre-culture and culture medium and presence of glucose (in our case cellobiose-born glucose) and can thus be better modulated by setting the suitable experimental conditions. Activity of β-glucosidase increases by the means of selection when *bglC*$^{+}$ *P. putida* is transferred from LB to M9 medium with cellobiose because this enzyme becomes essential for the cells. The activity of glucokinase increases even more because the expression of *glk* is probably very low in LB but is triggered in medium with cellobiose. De-repression of *glk* expression takes some time (Udaondo *et al*., 2018), therefore, the activity of glucokinase measured in cells pre-cultured in M9 medium with cellobiose (Fig. S4, M9-M9L) is the highest.

We believe that these results elucidate the observed accumulation of cellobiose-born glucose in the culture with *P. putida* EM42 pSEVA2213_*bglC* cells pre-grown in LB (Fig. 4A) and subsequent substantial reduction of free glucose concentration in the culture with the cells pre-grown in M9 medium with cellobiose (Fig. 4D).

### Supplementary tables

**Supplementary Table S1.** Strains and plasmids used in this study

| Strain or plasmid | Characteristics | Source or reference |
|---|---|---|
| ***Escherichia coli*** | | |
| Dh5α | Cloning host: F-λ-*endA1 glnX44*(AS) *thiE1 recA1 relA1 spoT1 gyrA96*(NalR) *rfbC1 deoR nupG* Φ80(*lacZ*Δ*M15*) Δ(*argF*-*lac*)*U169 hsdR17*($r_K^- m_K^+$) | Grant *et al*. (1990) |
| ***Pseudomonas putida*** | | |
| EM42 | Derivative of strain *P. putida* KT2440: Δprophages1,2,3,4 ΔTn7 Δ*endA1* Δ*endA2* Δ*hsdRMS* Δflagellum ΔTn*4652* | Martínez-García *et al*. (2014b) |
| EM42 Δ*gcd* | EM42 with scarless deletion of *gcd* gene (PP_PP1444) encoding periplasmic glucose dehydrogenase | Dvořák and de Lorenzo (2018) |
| **Plasmids** | | |
| pSEVA2213 | Expression vector: *oriV*(RK2) pEM7 *neo*, $Km^R$ | Silva-Rocha *et al*. (2013) |
| pSEVA2213_*bglC* | pSEVA2213 with *bglC* gene (SacI/PstI) | Dvořák and de Lorenzo (2018) |

### Supplementary figures

**Figure S1.** Recycling of *P. putida* EM42 resting cells converting D-xylose to D-xylonic acid.

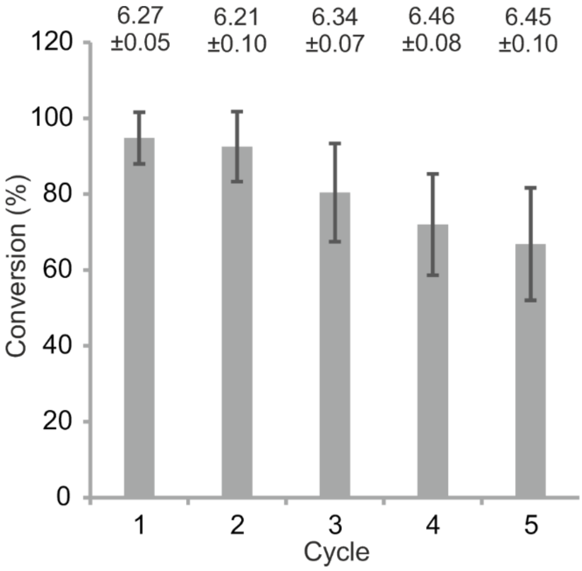


Medium with 5 g $L^{-1}$ D-xylose was inoculated using cells pre-culture in LB medium to the initial $A_{600}$ of 1.0. Cells were then incubated at 170 rpm and 30 °C. After 24 h, cells were centrifuged (4000 RPM, 15 min. at room temperature), pellets were re-suspended in fresh medium with xylose and supernatants were used for further quantifications. Xylose conversion to xylonate (g $g^{-1}$) was determined for each of the five cycles. The average $A_{600}$ determined at the beginning of cycles 1 - 5 was 0.99, 0.68, 0.51, 0.44, and 0.33, respectively. Columns represent means ± SD from at least four biological replicates from two independent experiments.

**Figure S2.** Two-day cultures of *P. putida* EM42 in minimal medium with 10 g $L^{-1}$ D-xylose and 10 g $L^{-1}$ D-glucose used as a sole carbon source for growth.

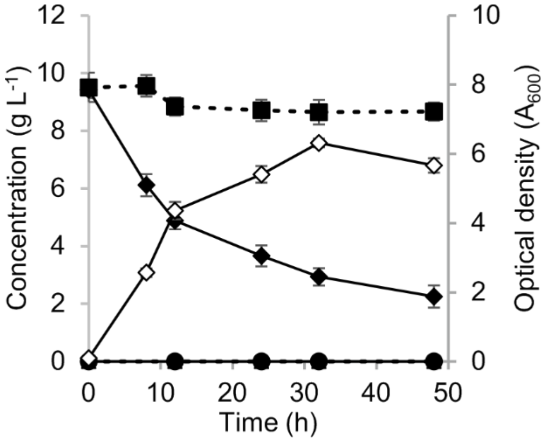


Experiment was carried out in 25 mL of minimal medium in flask shaken at 170 rpm and 30 °C. Minimal medium was inoculated to the initial $A_{600}$ of 0.1 using cells obtained from overnight culture in lysogeny broth. D-xylose, filled squares (■); D-xylonate, filled circles (●); D-xylono-λ-lactone, filled triangles (▲); D-glucose, filled diamonds (♦); cell biomass, open diamonds (◊). Data points shown as mean ± SD from three biological replicates.

**Figure S3.** Polyhydroxyalkanoate (PHA) accumulation in *P. putida* EM42 cells grown on diverse carbon sources including D-cellobiose.

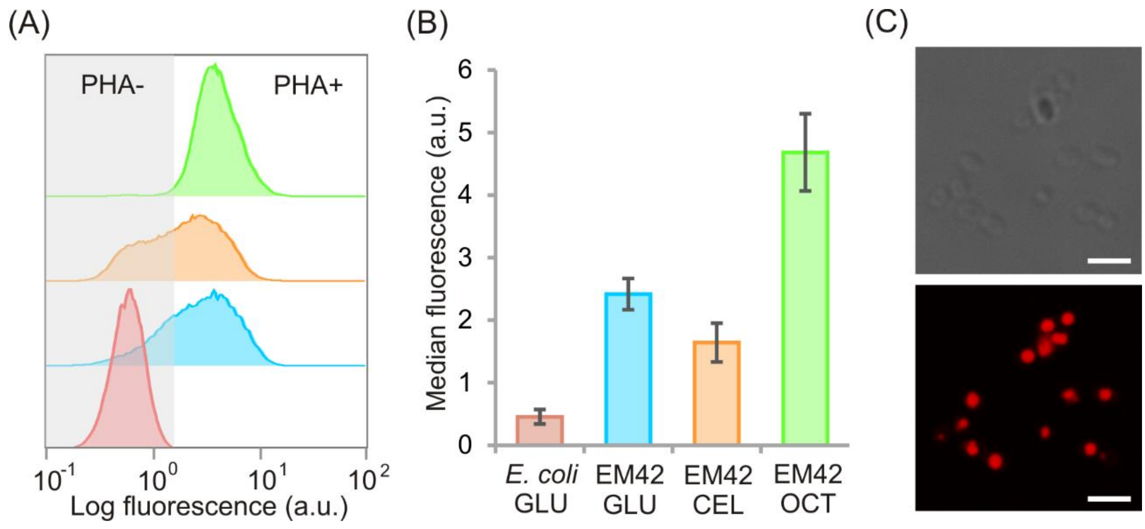


Cultures were carried out in 10 mL of nitrogen limited M9 minimal medium in flasks shaken at 170 rpm and 30 °C. (A) Representative flow cytometry histograms showing Nile red fluorescence of bacterial populations grown for 48 h with different carbon sources: red histogram, *E. coli* Dh5α pSEVA2213 grown on 30 g $L^{-1}$ D-glucose (*E. coli* GLU; negative control for PHA accumulation); blue histogram, *P. putida* EM42 pSEVA2213 grown on 30 g $L^{-1}$ D-glucose (EM42 GLU); orange histogram, *P. putida* EM42 pSEVA2213_*bglC* grown on 15 g $L^{-1}$ D-cellobiose (EM42 CEL); green histogram, *P. putida* EM42 pSEVA2213 grown on 20 mM octanoate (EM42 OCT; positive control for PHA accumulation). (B) Median fluorescence of bacterial populations measured by flow cytometry after 48 h of culture. Columns are mean ± standard deviation from three to four biological replicates. (C) Bright-field (upper photography) and confocal (lower photography) microscopy of EM42 CEL cells. White scale bar shows 2.5 µm distance.

**Figure S4.** Activities of β−glucosidase BglC (A) and glucokinase Glk (B) measured in cell-free extracts prepared from *bglC*$^{+}$ strain of *P. putida* EM42 grown in several different conditions.

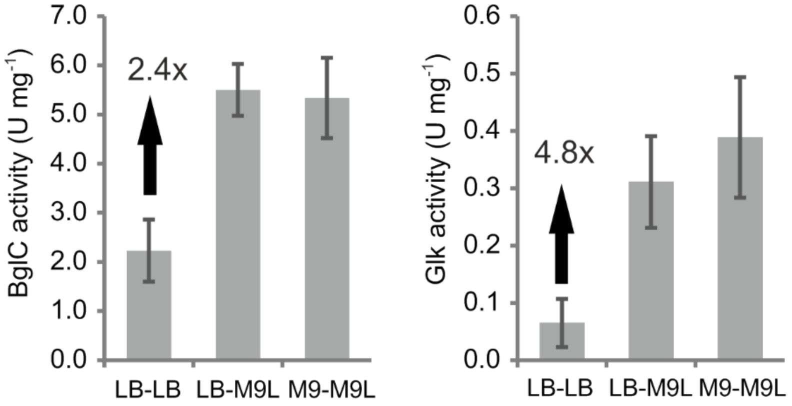


Cells were pre-grown overnight in lysogeny broth (LB) or in M9 minimal medium with standard nitrogen content (M9). Overnight cultures in LB were used to inoculate day cultures in LB (LB-LB) or in nitrogen-limited M9 medium (LB-M9L). Overnight cultures in M9 medium served for inoculation of day cultures in nitrogen-limited M9 medium (M9-M9L). When $A_{600}$ reached 0.5, day cultures were stopped and cell-free extracts prepared from the collected cells. Specific activities of BglC and Glk in extracts were measured as described in supplementary Experimental procedures. Black arrow with number shows X-times difference in activity of given enzyme in cells pre-grown and grown in LB (LB-LB) and cells pre-grown in LB and then grown in M9 medium with limited nitrogen content (LB-M9L). Columns represent means ± SD from three biological replicates and two technical replicates.
